# Multi-scale drivers of daily flow intermittency in a regulated desert river

**DOI:** 10.1101/2024.04.22.590594

**Authors:** Eliza. I. Gilbert, Thomas F. Turner, Melanie E. Moses, Alex J. Webster

## Abstract

Fluvial ecosystems are vital for biodiversity and human welfare but face increasing threats from flow intermittency caused by climate change and other human activities. To better understand drivers of flow intermittency, we analyzed long-term and spatially explicit river drying data from the Rio Grande, a regulated river in the North American desert southwest that was historically perennial but is now persistently intermittent. We examined the spatial structure and influences of precipitation, temperature, in-channel infrastructure, and river discharge on flow intermittency using multivariate autoregressive state space (MARSS) models and 12 years of daily data. Our findings indicate that river diversion rates at dams and irrigation return flows significantly structure the spatial occurrence of flow intermittency, but factors (possibly geologic) at distances ≤ 7 kilometers (km) are more influential as predictors of drying. Controlling influences of temperature and precipitation were not detected at the reach level (∼154 km) but were significant at each of the subreach scales (n = 3) investigated. At all subreach scales, temperature’s effect size exceeds precipitation by 2.5 times and is the strongest predictor of drying. Overall, process variance decreased by 98% between our reach- and all subreach models, suggesting that scale-sensitive models have great potential to accurately inform environmental flow management strategies aimed at mitigating negative effects of climate change and water extraction.

## 1. Introduction

Rivers and streams cover a small percentage of the Earth’s surface (∼2.3%) but are biodiversity hotspots where about 9.5% of the Earth’s animal species reside (Reid et al., 2019; Tockner et al., 2011). Human development has altered natural river flow and increased flow intermittency, causing significant concern because river and stream flow regulates key biogeochemical processes like nutrient cycling, provides essential habitat to resident biota adapted to these systems, and delivers ecosystem services to humans like access to surface water (Newbold 1992; Power et. al. 1995; Palmer and Ruhi 2019; Zipper et al., 2021). Naturally intermittent systems make up over half of the world’s river and streams, but the frequency and extent of flow intermittency is now intensifying in historically perennial systems (Datry et al., 2017). Intermittency effects are especially acute in arid regions affected by water resource development and climate change (Larned et al., 2010; Messager et al., 2021). Although it is understood that providing environmental flows could mitigate these trends, there remains a pressing need to identify drivers of flow intermittency to support actionable management options (Wineland et al., 2022; Gido et al. 2023).

Theory suggests flow intermittency is driven by three factors that function at different spatiotemporal scales (see Figure 5 in Costigan et al., 2016). These factors are meteorology, geology, and land cover, which are respectively proposed as first-, second-, and third-order controls. Meteorological controls scale down through four levels that begin with long-term climate patterns, to multi-annual variability, to inter and intra-annual variability, and ultimately to short-term discrete events such as a localized precipitation at the smallest scale. Geological controls are also hierarchically organized, scaling from the watershed to river reaches down to functional sets like a pool-riffle complex, and ending at an individual mesohabitat such as a single pool. Costigan et al. (2016) conceives of land cover as controlling flow intermittency through interactions between meteorology, geology, ecology, and human activities and this control scales broadly from biomes, to localized land use, to the riparian zone. At the smallest spatial scale, in-channel engineered structures like dams are forms of land cover that control flow intermittency.

Intuitively, dams should control flow intermittency because these structures interrupt the natural flow regime (Poff et al, 1997; Chalise et al., 2021). However, neither the impact of dams on downstream flow intermittency nor the direction of influence are consistently discernible (Costigan et al., 2016; Belemtougri et al. 2021). Obtaining a clearer understanding of the impact of in-channel engineered structures such as large dams or smaller irrigation diversion dams is important because approximately 2.8 million dams regulate river flows worldwide, and many more are proposed (Lehner et al, 2011; Grill et at., 2015). Within the continental United States more than one-third (1.2 million miles) of rivers and streams are regulated by such structures (Carlisle et al., 2019).

This study is motivated by the need to better understand the impacts of dams and other in-channel infrastructure on flow intermittency in a backdrop of meteorological, geological and land cover controls. We focused on the reach and subreach scales where in-channel engineered structures such as dams are most likely to control flow intermittency (Costigan et al., 2016). We used Costigan et al.’s (2016) definition of a reach as a length of river composed of integrated geologic units with similar characteristics of valley confinement and channel sinuosity (Costigan et al., 2016), and defined the subreach scale at varying extents that are less than the reach scale.

A significant challenge in understanding the mechanisms driving flow intermittency lies in the potential for the effect of controls to vary across spatial scales (Costigan et al., 2016). For example, at a continental scale, flow intermittency regimes of 894 gaged rivers and streams are more influenced by land cover attributes than by meteorology or geology (Price et al., 2021). In contrast, watershed-scale studies suggest that geologic controls, such as catchment area, profile curvature, soil hydraulic conductivity, and bedrock permeability, are key predictors of spatiotemporal flow intermittency rather than land cover (Kaplan et al., 2022). These empirical studies underscore the difficulty in understanding flow intermittency and the potential for larger or smaller-scale controls to dampen on another’s effect (Fritz et al., 2020; Brown et al., 2023). Thus, we also designed our study to assess drivers of flow intermittency across spatial scales.

We formulated two questions to examine drivers of flow intermittency. First, we asked what factors structure the spatial pattern of daily flow intermittency? We hypothesized that modeling flow intermittency at spatial scales related to in-channel infrastructure would more accurately predict daily dynamics compared with models that focus solely on either the broadest reach or smallest subreach scale. This hypothesis is based on the premise that the location of surface water extraction and return flow regulated by in-channel infrastructure is a key determinate of the spatial structure of flow intermittency. Second, we asked what are the relative contributions of proximate measures of precipitation and temperature, in-channel infrastructure, and within reach discharge to daily flow intermittency across spatial scales? We predicted that precipitation and temperature predominately drive flow intermittency at the subreach scale, and the influence of in-channel infrastructure becomes more pronounced as subreach size decreases. Here, river discharge provides insight into the cross-scale interaction between meteorology and land cover because it reflects the influence of snowmelt runoff driven by climate and smaller-scale human constraints on flow arising from upstream regulation of surface flow. We show that a deeper understanding of flow intermittency drivers across spatial scales can enhance our ability to conserve, manage, and protect freshwater ecosystems by providing targeted options for environmental flow (Costigan 2016).

## 2.0 Methods

### 2.1 Study area

Our study focuses on the Rio Grande because contemporary flow intermittency in this large and historically perennial North American desert river exemplifies the global challenges faced in sustaining freshwater ecosystem functions and services (Garza-Díaz and Sandoval-Solis 2022). The Rio Grande is contained within the Rio Grande/Rio Bravo Basin (Fig. 1), which encompasses approximately 11% of the continental United States (Sandoval-Solis et al., 2022). The Rio Grande is the basin’s largest river and is noteworthy as the fifth-longest river in the United States and twentieth longest worldwide (Patiño-Gomez et al., 2007). While droughts in this snowmelt-driven system may have occasionally caused flow intermittency, modern water management practices combined with climate change have intensified flow intermittency, making it a persistent occurrence (Garza-Diaz & Sandoval-Solis, 2022; Wineland et al., 2022). Nearly 83% of the basin’s water resources are diverted for agriculture, suggesting that understanding how in-channel infrastructure like diversion dams control flow intermittency would contribute to efforts to develop environmental flows in this and other arid agricultural systems (Sandoval-Solis et al., 2022)

**Fig. 1.**
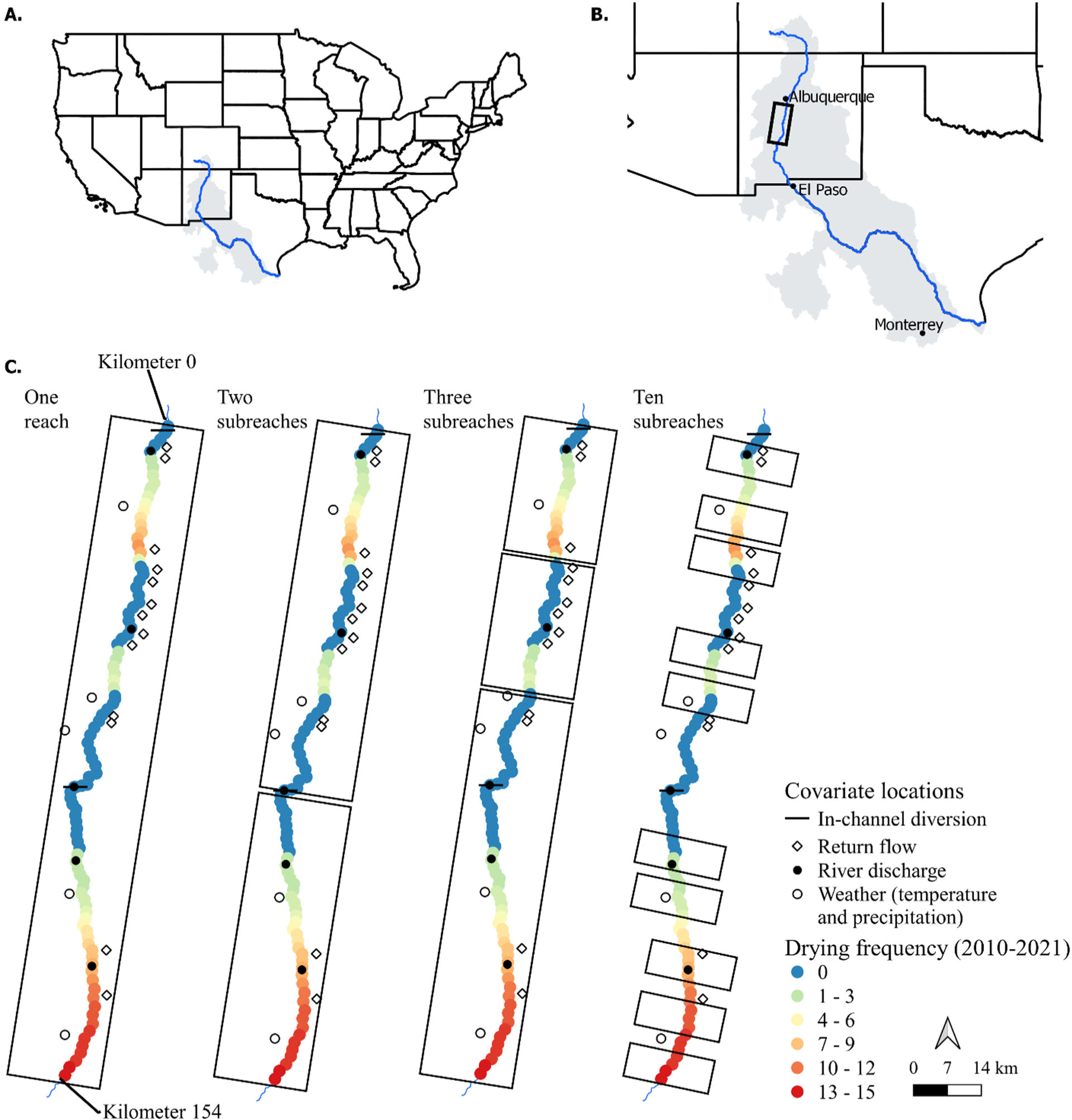
Panel A, the Rio Grande/Rio Bravo Basin (shaded in grey) and the Rio Grande (blue line with flow north to south). Panel B, the location of study reach (black box). Panel C, spatial structuring of the models used to assess daily flow intermittency spatial structure and controls.

Significantly, the Rio Grande is home to the endangered Rio Grande Silvery Minnow (*Hybognathus amarus*). When the river is diverted to provide crop irrigation, close monitoring of flow intermittency is required to support the fish’s conservation (Platania et al., 2020; McKenna, 2021; Walsworth and Budy, 2021). Accordingly, there are consistent annual flow intermittency observations over a ∼154-kilometer (km) reach of the Rio Grande that supports the minnow and we used this river reach as our study area (Fig. 1). Monitoring of flow intermittency in this reach of the Rio Grande was conducted at an unprecedented temporal and spatial resolution with daily observations made approximately every 0.16 km (i.e., a tenth of a mile; McKenna et al., 2023). Such long-term and detailed daily flow intermittency data are rare (Sabo, 2014; Allen et al., 2019; Durighetto et al., 2020), offering a unique opportunity to analyze the spatial dynamics of flow intermittency in this regulated river system (Garza-Diaz and Sandoval-Solis 2022).

### 2.2 Flow intermittency metric and covariates

We evaluated our hypotheses using daily time-series data restricted to the irrigation season when flow intermittency occurs (April to October) and calculated flow intermittency as the daily cumulative km-days dry. We chose this metric to characterize flow intermittency because annual cumulative km-days dry is a recognized hydrologic driver of yearly Rio Grande Silvery Minnow abundance (Walsworth and Budy, 2021). We calculated daily cumulative km-days dry by adding the number of dry km in a day to the cumulative total from the prior day. The strength of this metric lies in its ability to capture both the spatial and temporal extent of flow intermittency (Palmer and Ruhi, 2019). We considered using the daily spatial extent of channel drying or its rate of change, but rejected these metrics because they violated normality and independence assumptions that underpin linear regression. Ultimately, we calculated daily cumulative km-days dry for 12 years (2010-2021) of data obtained from the River Eyes Monitoring Program online portal (GSA 2023).

To assess individual and combined effects of meteorology and land cover on daily flow intermittency, we collected publicly available data from measurement stations within our study area (Fig. 1). We obtained temperature (°C) and precipitation (mm) from five locations with records sourced from the National Centers for Environmental Information (NOAA 2023). In our study area, in-channel infrastructure consists of two cross-river diversion dams (Supplemental Fig. S1) and 13 canals that return unused irrigation water to the Rio Grande. We accessed data on the rate of diversion (m^3^/s) at each dam and rate of return (m^3^/s) for each canal from the Middle Rio Grande Water Management Toolbox (USBOR 2023). To elucidate the interaction between larger scales meteorologic variability and in-channel infrastructure management, we obtained upstream discharge (m^3^/s) for each subreach using five gages (USGS 2023). Although there were more than five discharge gages in the study area, we restricted our use to those that consistently acquired data from 2010-2021 because this was the interval we used to calculate our flow intermittency metric.

### 2.3 Modelling framework

We employed the multivariate autoregressive state-space (MARSS) modeling framework to evaluate the spatial structure of daily flow intermittency and to quantify its drivers. We selected MARSS for its robust capacity in elucidating temporal variability across multiple time series response variables, owing to its ability to effectively handle temporal autocorrelation. The core of MARSS involves matrix calculations, which allow an explicit evaluation of spatial relationships (Jankowski et al., 2021; Holmes et al., 2023; Supplemental Fig. S2). Furthermore, MARSS provides the advantage of distinguishing between process variance caused by environmental stochasticity and variation stemming from observational error, thus enabling a more precise interpretation of the environmental factors driving daily flow intermittency compared to other types of linear regression.

MARSS partitions variance from observational error using a two-component model. The first component, the process model (Eq. 1), describes changes over time in the unobserved but true process states of nature (*x*_*t*_) and the effect of covariates (*C*_*t*_*c*_*t*_) on these states (Holmes et al. 2023). The *C*_*t*_ matrix represents the parameters for each covariate measured at time *t* and maps their effects onto the process states. The *c*_*t*_ vector contains each of the measured covariates appropriately scaled to the spatial configuration of each process state. The second model component, the observation model (Eq. 2), relates the observed data (*y*_*t*_) to the process states (*x*_*t*_).

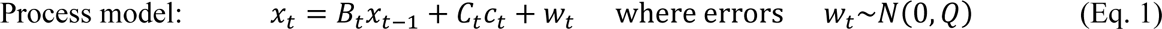

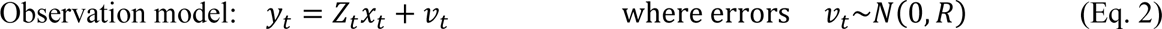

Through design of the *Z*_*t*_ matrix, the user specifies the number of process states and relationship among these states (e.g., shared or independent temporal variation). This can occur because the *Z*_*t*_ matrix acts to map each observed time series onto the unobserved states estimated in the process model as *x*_*t*_. Within the process model, the vector *w*_*t*_ contains process variance distributed as multivariate normal with a mean vector 0 and covariance matrix *Q*. The diagonal terms of the *Q* matrix can be specified by the user to represent different process variance such as those that are equal or unequal across states. The off diagonal of the *Q* matrix provides another estimate of shared behavior across states because these terms represent covariance in process variance after accounting for covariate effects (Jankowski et al., 2021). Thus, using the flexibility provided in the *Z*_*t*_ and *Q* matrices, we constructed models with hypothesized spatial structures and process variance (Supplemental Fig. S2).

Other model parameters include the *B*_*t*_ matrix that determines the degree to which each unobserved process state reverts to its mean. The vector *v*_*t*_, contains observation errors for each observed times series. As with the *w*_*t*_, it is multivariate normal with a mean vector of 0 but has a covariance matrix *R*. In our models, we specified equal parameter estimates for *B*_*t*_ and *R* to respectively estimate mean reversion and observation error uniformly across times series. For each process state, we set the initial condition of *x* to time = 0.

### 2.4 Spatially-structured models and covariates

To test our hypothesis regarding the spatial structure of daily flow intermittency, we developed four MARSS models, each predicting flow intermittency at increasingly smaller spatial scales (Fig. 1). The largest spatially-scaled model described temporal variance in flow intermittency as a single reach bounded at the top by a diversion dam and at the bottom by the most downstream consistent occurrence of flow intermittency (154 km downstream). Hereafter, we refer to this as the one-reach model. We introduced a smaller-scale two-subreach model whose subreaches were bounded by the upstream locations of cross-river diversion dams. The upstream subreach was 86 km and the downstream subreach was 68 km long. Visual inspection of flow intermittency over the 12-year dataset suggested effects potentially related to the location and density of irrigation return canals. This observation led to the construction of a third model at an even smaller spatial scale, consisting of three subreaches. These subreaches were delineated based on observable changes in flow intermittency frequency over the 12-year period, and we demarcated the subreach by the highest frequency of flow intermittency and upstream points of flow intermittency reemergence (Fig 1c). We refer to this model as the three-subreach model and moving from upstream to downstream, the subreaches measure 32, 32, and 90 km in length.

Models were constructed at the smallest spatial scale by randomly generating ten non-overlapping 7-km subreaches and repeating this process 100 times to ensure comprehensive sampling of the study area (Fig. 1). The rationale was based on findings by Yackulic et al. (2022), who estimated that a 10% increase in wetted river reach length—equivalent to 7 km — would increase annual Rio Grande Silvery Minnow abundance by a commensurate 10%. Due to infrequent flow intermittency in some reaches, 45 models had poor convergence, resulting in 55 converged models. We hereafter refer to these 55 models as the ten-subreach models.

For each model, we calculated daily cumulative km-days dry at both the reach and subreach scales. The spatial nature of our study required covariates be spatially explicit, reflecting the distinct characteristics of each subreach (Table 1). For example, we obtained daily discharge from the nearest gauge upstream of each subreach. Additionally, to align with our calculation of cumulative km-days dry, we also aggregated our covariates cumulatively, ensuring that the analysis reflects the influence of these drivers over time (Adams et al., 2020; Szeitz and Moore 2022). Prior to model computation, we conducted Pearson’s correlation analysis of covariates to detect strong linear relationships. Pairwise correlation coefficients were at or below 0.28 (indicating low to moderate intercorrelation) and we included all five covariates (temperature, precipitation, diversion rate, return flow rate, and upstream discharge) in regression models. To normalize each flow intermittency and covariate time series, we applied z-score transformations prior to model computation.

**Table 1.** Covariate calculations for each spatially scaled model. Values in parentheses are the number of gages and “upper”, “middle”, or “lower” indicates the relative location of subreaches.

| Covariate | One reach | Two subreaches | Three subreaches | Ten subreaches |
| --- | --- | --- | --- | --- |
| Precipitation | Sum (5) | Sum (upper = 3)<br>Sum (lower = 2) | Sum (upper = 1)<br>Sum (middle = 2)<br>Sum (lower = 2) | Nearest |
| Temperature | Mean (5) | Mean (upper = 3)<br>Mean (lower = 2) | Mean (upper = 1)<br>Mean (middle = 2)<br>Mean (lower = 2) | Nearest |
| Diversion | Mean (2) | Nearest upstream (1) | Nearest upstream (1) | Nearest upstream |
| Return flow | Sum (13) | Sum (upper = 11)<br>Sum (lower = 2) | Sum (upper = 3)<br>Sum (middle = 6)<br>Sum (lower = 4) | Sum of upstream |
| River discharge | Mean (5) | Nearest upstream (1) | Nearest upstream | Nearest upstream |

### 2.5 Model evaluation and parameter estimates

We used root mean square error (RMSE) to assess the performance of four spatially-scaled MARSS models and only compared models whose process states were specified as independent (i.e., *Z*_*t*_ matrix diagonals = 1; Supplemental Fig. S2). We also explored models with shared process states (i.e., *Z*_*t*_ matrix with columns whose rows shared 1s; Supplemental Fig. S2) but excluded these from further analysis due to non-normally distributed and temporally autocorrelated errors. For the two-, three-, and ten-subreach models, we used Akaike Information Criterion for small samples sizes (AICc) to identify the most suitable structure to relate process variance across subreaches (i.e., Q matrix structure). We compared three Q matrix (i.e., covariance) structures where we specified the diagonal as equal, unequal, or unconstrained variance and covariance. For the ten-subreach models, we randomly selected three time series to assess the most suitable Q matrix (Supplemental Fig. S2). From the AICc results, we used the best fitting Q matrix for each spatially-defined model to calculate parameters estimates.

We refined parameter estimates using the Broyden-Fletch-Goldfarb-Shanno (*BFGS*) maximum-likelihood algorithm, with initial parameters informed by models calculated from the expectation maximization (*EM*) likelihood algorithm (Holmes et al., 2023). For the reach model and best performing subreach models, we obtained parameter means and 95% confidence intervals (CIs) using the *MARSSparamsCIs* function. For the ten-subreach model, we used a bootstrap method and 1,000 replicates to calculate an overall mean and 95% CI from the 55 estimated parameter means (Efron and Tibshirani 1986). We considered the effect of a covariate to be insignificant if 95% CIs encompassed zero and considered the effect of a covariate to be similar across models when 95% CIs overlapped. All modeling was conducted in the R statistical environment using the *MARSS* package (R Core Team 2021; Holmes et al., 2023).

## 3.0 Results

### 3.1 Flow intermittency and covariate summaries

Flow intermittency in our study area was highly variable with annual values varying from 398 to 6,915 cumulative km-days dry (mean = 2,560; sd = 2,256) and differential subreach contribution (Fig. 2 and 3). Temperature and precipitation conditions were more stable. Over the 225 days of the irrigation season, annual cumulative temperatures spanned a narrow range from 6,355 to 6,769 °C (mean = 6,541; sd = 138). However, precipitation was more varied and ranged from 482 to 1,142 mm (mean = 820; sd = 218).

**Fig. 2.**
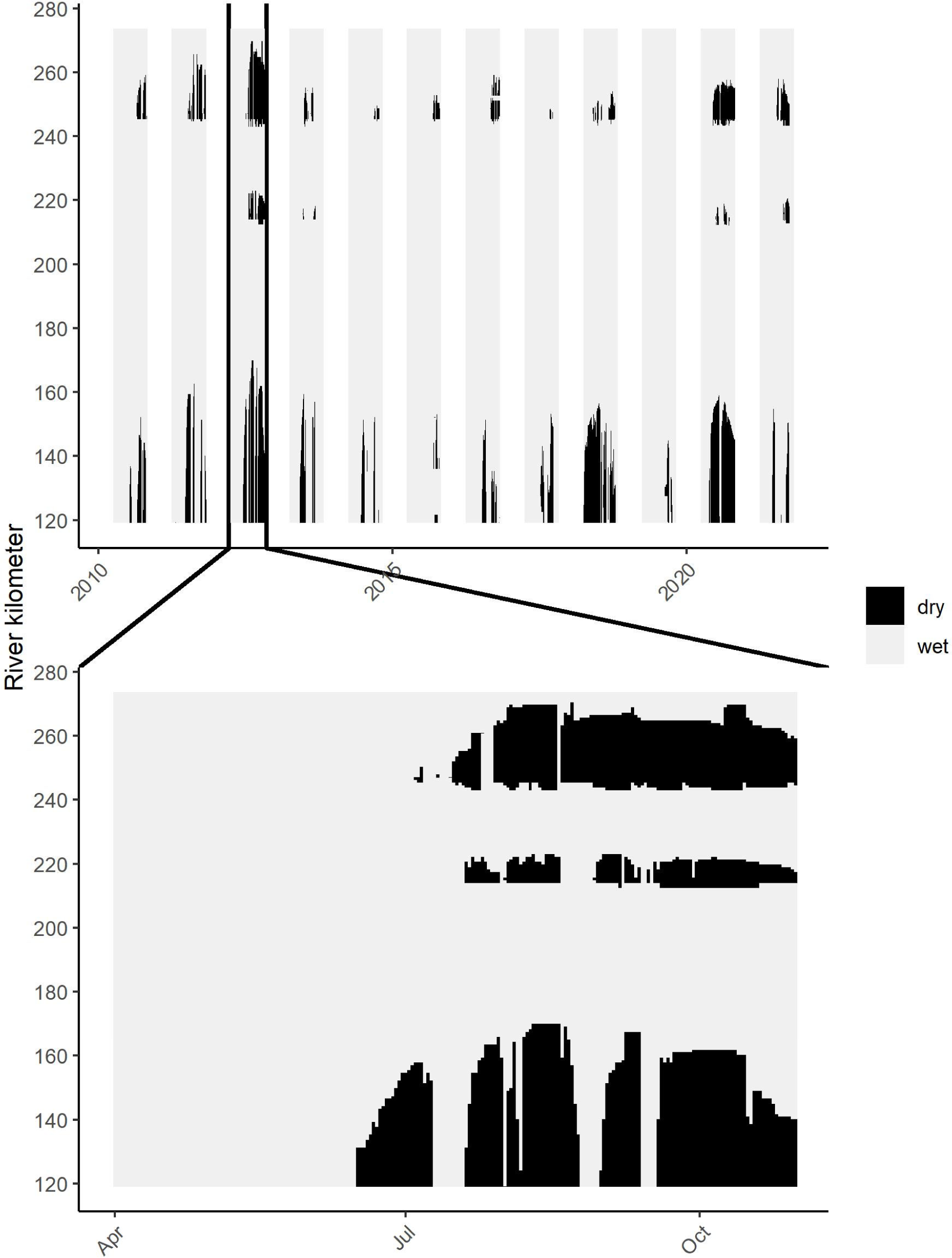
Daily flow intermittency spatial and temporal distribution in the study area during each irrigation season (Apr – October). Bottom panel depicts 2012.

**Fig. 3.**
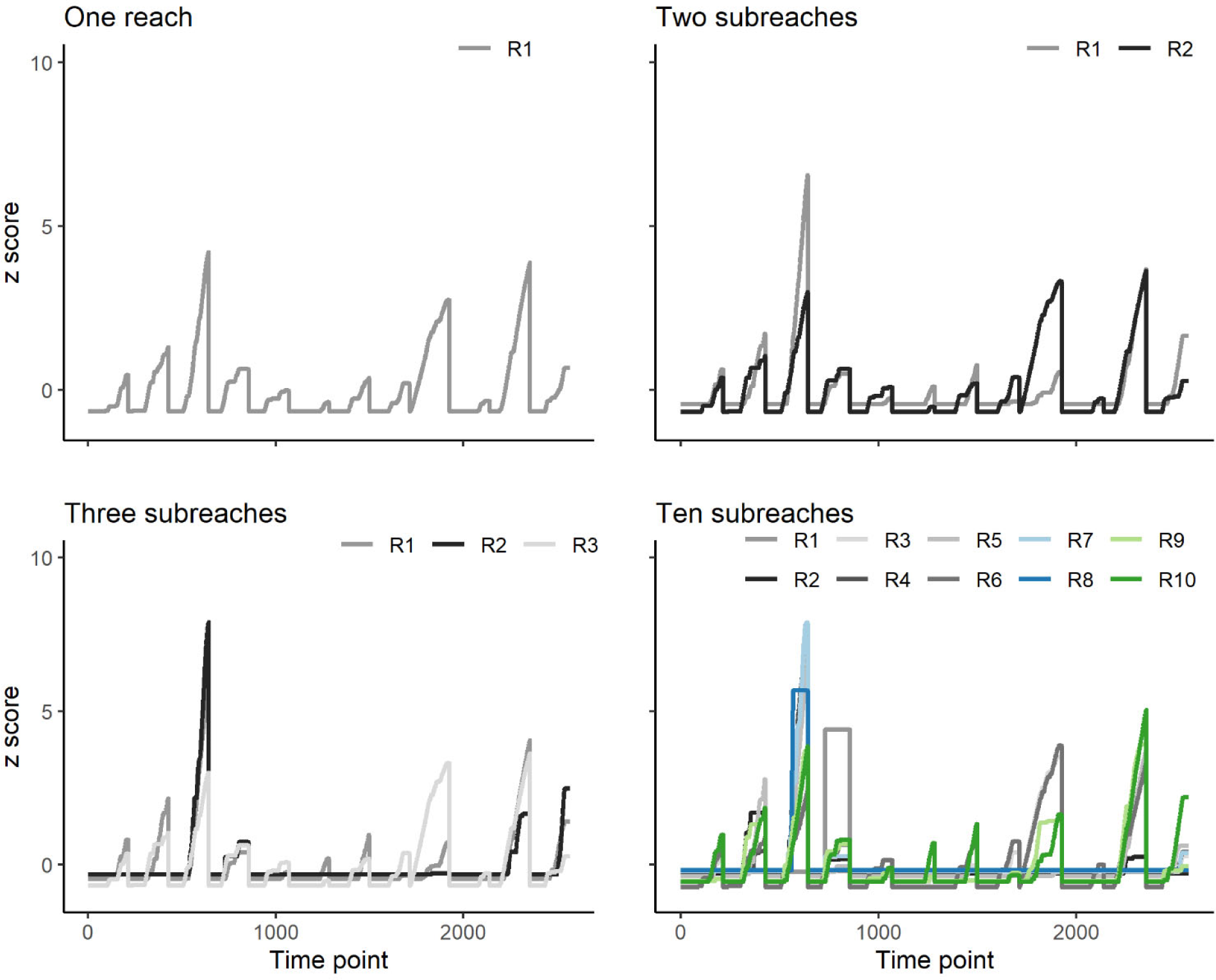
Daily time series of cumulative kilometer-days dry (2010 to 2021) for each spatially scaled model.

The two in-channel diversion dams showed considerable annual variability in their operations. On a daily basis, they diverted anywhere from 0 to 100% of instream flow and, on average, the upstream diversion diverted 39% of flow while the downstream diversion diverted 23%. Annual diversions ranged from 1,186 to 4,847 m^3^/s each year (mean = 2,687; sd = 770 m^3^/s). Approximately 34-45% of diverted water was returned to the river, equating to annual cumulative returns ranging from 648 to 2,103 m^3^/s (mean = 1,145; sd = 438 m^3^/s). At any one irrigation return canal, annual daily flow ranged from 0 to 6 m^3^/s and the largest annual mean return from a single canal was 4 m^3^/s. Despite the variability in river diversion and return flow rates, average cumulative discharge in the study area was consistent, ranging from 4,971 to 4,987 m^3^/s (mean = 4,975; sd = 7).

### 3.2 Flow intermittency spatial structure

Across all models, our MARSS models predicted flow intermittency with high accuracy. This was reflected by RMSEs approaching zero (range = 4.8 x 10^-17^ to 2.1 x 10^-10^; Table 2). For the two- and three-subreach models, unconstrained process variance and covariance among process states yielded the best fitting models (Δ AICc ≥ 4; Table 3). For the ten-reach models, unequal process variance provided the best-fitting model compared to equal process variance (Δ AICc ≥ 4), but we were not able to consider unconstrained process variance and covariance due to insufficient data. Observation error estimates were negligible across all models (range = 1.96 x 10^-14^ to 2.13 x 10^-11^). The highest process variance was calculated for the one-reach model (mean = 1.02; 95% CI = 0.93 to 1.12), while process variance for the smaller spatially-scaled models was similar among models with each estimated mean equal to 0.02. Mean reversion rates were consistent across all models (range = 0.98 to 1.00).

**Table 2.** Number of multivariate autoregressive state space model process states in each *Z*_*t*_ matrix used to assess daily flow intermittency spatial structuring (Supplemental Fig. S2). The number of process states is equal to the number of reaches (one) or subreaches (two, three, or ten) delineated within the study area. Model fits are reported as root mean square error (RMSE).

| Number of<br>process states | Relationship among<br>process states | RMSE |
| --- | --- | --- |
| One | Not applicable | $4.8 \times 10^{-17}$ |
| Two | Independent | $9.2 \times 10^{-17}$ |
| Three | Independent | $1.1 \times 10^{-14}$ |
| Ten | Independent | $2.1 \times 10^{-10}$ |

**Table 3.** Comparisons of multivariate autoregressive state space process variance (i.e., Q matrix) comparisons for each spatially scaled model predicting the daily accumulation of dry kilometers. The number of process states is equal to the number of subreaches delineated within the study area.

| Process states | Process variance | Estimated parameters | Iterations for convergence | Log-likelihood | $\Delta AICc$ |
| --- | --- | --- | --- | --- | --- |
| Two | Unconstrained | 11 | 24 | 3719 | 0 |
|  | Unequal | 10 | 16 | 2259 | 2918 |
|  | Equal | 9 | 17 | 2227 | 2980 |
| Three | Unconstrained | 14 | 25 | 6701 | 0 |
|  | Unequal | 11 | 19 | 3161 | 7074 |
|  | Equal | 9 | 16 | 3123 | 7146 |
| Ten * | Unequal | 18 | 24 | 12631 | 0 |
|  | Equal | 9 | 15 | 12324 | 595 |
|  | Unconstrained | - | - | - | - |

\*Log-likelihood and $\Delta AICc$ for these models is the mean of three models randomly chosen from the 55 computed models. An unconstrained (variance and covariance) $Q$ matrix was not considered due to insufficient data.

### 3.3 Covariate parameter estimates across spatial scales

Results from the one-reach model suggested temperature, precipitation, in-channel infrastructure, and discharge were not significant predictors of daily flow intermittency (Fig. 4). In contrast, these covariates influenced flow intermittency in the two-, three-, and ten-subreach models, with temperature and precipitation exhibiting similar effects sizes and direction across all subreach models. With a positive relationship to flow intermittency, temperature had the most pronounced impact with mean estimates ranging from 0.024 to 0.034 and differences among smaller-scaled models were insignificant (Supplemental Table S1). Precipitation displayed a consistent negative relationship with flow intermittency at subreach scales. Mean estimates ranged from -0.009 to -0.011 and the impact of precipitation was also similar across all subreach models (Supplemental Table S1).

**Fig. 4.**
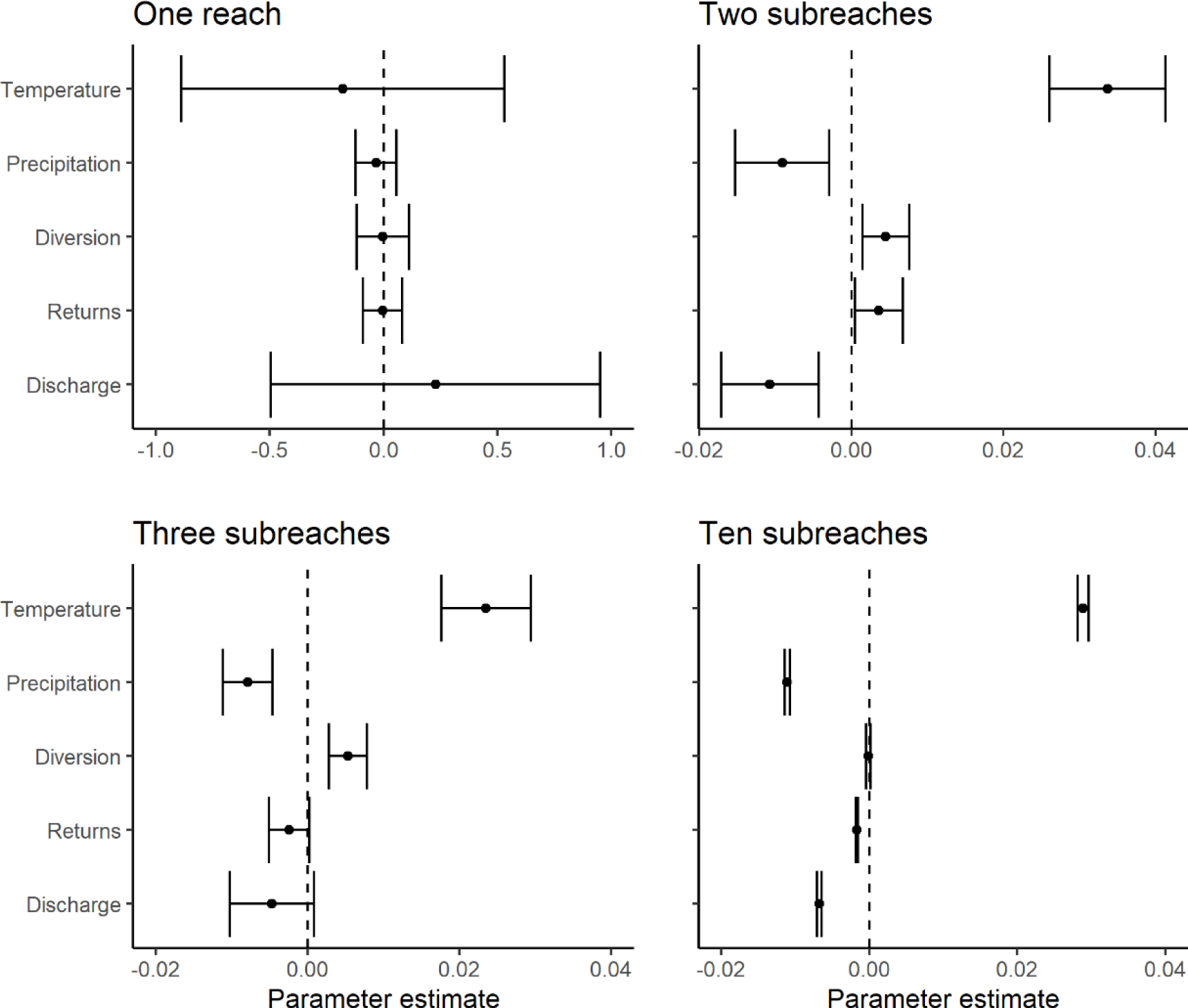
Covariate parameter estimates (mean and 95% confidence intervals) predicting daily flow intermittency across spatially structured models. All x-axes are on the same scale, except for the one-reach model.

In contrast, in-channel infrastructure’s relative impact on flow intermittency varied across subreach models. At the two- and three-subreach scales, irrigation diversions were positively correlated with flow intermittency with similar mean estimates of 0.005 (95% CI = 0.001 to 0.008 and 0.003 to 0.008, respectively). Yet, diversion rates were not a significant factor at the ten-subreach scale (Fig. 4). The two-subreach model suggested a modest correlation between the rate of diversion and returns, as returns also demonstrated a positive relationship to flow intermittency (mean = 0.004; 95% CI = 0.000 to 0.006). Returns did not significantly influence flow intermittency at the three-subreach scale and exhibited a negative correlation at the ten-subreach scale (mean = -0.002; 95% CI = -0.002 to -0.001).

In general, river discharge was negatively associated with flow intermittency at subreach scales. Both the two- and ten-subreach models demonstrated a negative relationship, with mean estimates of -0.011 (95% CI = -0.017 to -0.004) and -0.008 (95% CI = -0.007 to -0.006), respectively. Although the mean discharge estimate for the three-subreach model also tended towards negative, it was not significant (-0.0050, 95% CI = -0.010 to 0.001).

## 4.0 Discussion

### 4.1 Factors structuring the spatial pattern of subreach daily flow intermittency

A key outcome of our study was the substantial reduction (98%) in process variance when transitioning from the reach to the subreach models. This reduction suggests that modeling at the subreach scale incorporates significantly more environmental stochasticity compared to broader scales. Consequently, integrating subreach spatial controls in future numerical or process-based hydrology models could enhance our understanding of flow connectivity and intermittency (Jaeger et al., 2019; Prancevic and Kirchner, 2019; Price et al., 2021). While our study points to the importance of small-scale geologic drivers in structuring daily flow intermittency, the acquisition and utilization of such detailed data can pose significant challenges (Horton et al., 2021). Our findings suggest that reliance on intricate data and hydrological models may not be required, as we demonstrate that incorporating universal meteorologic controls like temperature and precipitation with smaller scale land cover attributes can effectively predict daily flow intermittency.

We found that while in-channel engineered infrastructure played a significant role in spatially structuring flow intermittency, factors at scales smaller than 7 km also strongly influenced its location. This inference is drawn from our two and three-subreach models, where the best-fitting models indicated disparate process variance across subreaches, suggesting that flow variation was best explained at subreach scales (Smits et al., 2019). Even at the smallest spatial scale (i.e., ten-subreaches), the best performing model had unequal process variance, indicating factors influencing the spatial structure of flow intermittency at 7 km were dissimilar among subreaches.

This finding necessitates a reevaluation of our initial hypothesis. We thought the sandy, alluvial nature of our study area would result in general homogeneity at the smallest geologic scale (i.e., grain size). We expected such homogeneity would render small-scale geologic drivers less influential in spatially structuring daily flow intermittency compared variability from in-channel infrastructure operations (i.e., dams), despite others classifying infrastructure as a third-order control relative to geology’s second-order influence (Costigan et al., 2016; Pearce and Kelson, 2004). However, our ten-subreach model suggests geologic heterogeneity at distances ≤ 7 km. The influence of geologic heterogeneity on flow intermittency can result from differences in sediment size, which regulate infiltration rates and thus, transmission loss (i.e., surface water loss to subsurface flow) (Fritz et al., 2020; Jacobs et al., 2020). Additionally, variable transmission loss due to sediment size can interact with the distance to groundwater and evapotranspiration (Soylu et al. 2011). Therefore, our results indicate that small-scale geologic heterogeneity, alongside other geologic and land cover factors, could play a crucial role in structuring daily flow intermittency, potentially overshadowing the influence of in-channel infrastructure.

### 4.2 Relative contributions of daily flow intermittency drivers across spatial scales

Our results corroborate the hypothesis that temperature and precipitation, more than in-channel infrastructure or proximate upstream discharge, is the primary driver of daily flow intermittency at the subreach scale. Although our analysis was focused on the reach and subreach scales and did not address larger scale impacts such as those from increased upstream water availability, we observed consistent influences of both temperature and precipitation on flow intermittency across all subreach scales, despite the potential impact from spatially heterogeneous transmission loss. This finding diverges from other subreach-scale studies whose results suggested either precipitation or temperature as singular drivers of flow intermittency (Yu et al., 2018; Moidu et al., 2021). Significantly, temperature showed a 2.5 times higher impact on flow intermittency than precipitation, contrasting with studies in headwater and mountain streams where precipitation is the predominant driver (Ward et al., 2018; Durighetto et al., 2020). These insights may indicate that, in arid conditions, temperature plays a more critical role in regulating subreach daily flow intermittency. As climate change progresses, we propose that temperature will play an increasing role in regulating daily flow intermittency (Zarch et al., 2017).

Our findings did not support our secondary hypothesis, which posited a more pronounced influence of in-channel infrastructure at smaller scales. Instead, our analysis suggested dynamic in-channel infrastructure controls of flow intermittency across spatial scales, as posited by Costigan et al. (2016). At the broadest subreach scale, diversions and flow returns similarly affected flow intermittency. However, this pattern shifted at smaller scales in which the directional influence of flow returns reversed, while the impact of diversion dams diminished and ultimately became undetectable. This result suggests that detecting the influence of in-channel infrastructure on flow intermittency or other flow regime metrics may be challenging at large and very small spatial scales (Chalise et al., 2021; Brown et al., 2023). This challenge may result from the scale of measurement or actual shifts in drivers at large and very small scales (Smits et al., 2019; Fritz et al., 2020). Addressing these scale extremes in future research would advance our understanding of scale-dependent drivers of flow intermittency.

### 4.3 Flow intermittency spatial management and modeling

The Rio Grande, identified as one of the world’s most at-risk rivers, requires specialized environmental flow strategies (Wong et al., 2007). Some have suggested that these strategies should focus on agricultural water use hotspots rather than adopting a basin-wide approach (Wineland et al., 2022). Our study, conducted in such a hotspot of high agricultural water use, offers water managers actionable insights to reduce flow intermittency. From our models, we estimate that increasing discharge by approximately 100 m^3^/s at the two-subreach scale, coupled with a corresponding decrease in diversions, could reduce daily flow intermittency accumulation by ∼1.52 km. At 7-km intervals, our results suggest this strategy would reduce accumulated flow intermittency by ∼1 km per day. Our findings suggest a nuanced approach could be developed to optimize water releases based on temperature variations to benefit the Rio Grande ecosystem.

Calculating flow intermittency as a daily accumulation introduces a level of abstraction that can be challenging to interpret. It is inherently difficult to model intermittency based on simpler metrics, such as the daily extent and change in flow intermittency, because of rapid shifts between flow and non-flow states (Kaplan et al., 2022). The threshold between these states is dynamic (Jacobs et at., 2020) making these simpler metrics difficult to predict. We recommend that future research focus on exploring the mathematical nature of these simpler daily flow intermittency metrics in a time series to identify tractable modeling frameworks (Huffaker et al., 2017).

### 4.4 A broader understanding of flow intermittency drivers

Overall, results from this study resonate with the well-established concept that physical and ecological processes function differently across spatial scales (Hefferenan et al., 2014; Fritz et al., 2020). At one end of the spectrum, our study found that temperature and precipitation, in-channel infrastructure, and instream discharge did not control flow intermittency at a 154-km-long reach scale. In contrast, at our smallest subreach scale, the specific influence of diversions was undetectable. These observations highlight that results of flow intermittency studies are closely tied to the spatial scale of analysis. This perspective could shed light on the varied findings in global, continental, and watershed analyses regarding the effects of dams and land cover on flow regimes and intermittency (Kaplan et al., 2020; Belemtougri et al. 2021; Chalise et al., 2021; Price et al., 2021; Brown et al., 2023).

Our study further emphasizes the need to analyze the interplay between social and ecological factors for sustainable water management (Sandoval-Solis et al. 2022). By investigating the combined effects of in-channel infrastructure, temperature, precipitation, and discharge across various spatial scales, our research provides site-specific and actionable solutions to addressing this challenge. For example, we calculated how much 100 m^3^/s decrease in diversion, increase in discharge, or increase in returns would reduce daily accumulated flow intermittency at the two- and ten-subreach scales. Importantly, these finding can contribute to development of options that can bridge the gap between providing environmental flows based on the natural flow regime and implementing emerging concepts like “designer flows” that are being developed to incorporate the realities of anthropogenically altered freshwater ecosystems and human needs (Poff et al., 1997; Poff 2018; Tonkin et al., 2021; Patterson et al., 2022).

### 4.5 Conclusion

Given the global scale of human alterations to the water cycle, it has never been more important to understand how climatic, geomorphologic, biological, and anthropogenic factors interact to mediate the structure and function of rivers (Brown et al., 2023). Our study responds to this urgent need by examining the spatial dynamics and determinants of daily flow intermittency in the Rio Grande at both a reach and subreach scales. Our findings highlight the significant influence of in-channel infrastructure on flow patterns but align with theoretical predictions that identify small-scale hydrogeologic factors as a more primary control. The research conducted here enhances our understanding of site-specific variables influencing flow intermittency and identifies the crucial role of spatial scale in flow intermittency analyses. We consider this finding as vital to incorporate into future analyses whose purposes are to manage flow intermittency in endangered river and stream ecosystems (Thompson et al., 2011; Fritz et al., 2020).

## CRediT authorship contribution statement

Eliza I. Gilbert: Conceptualization, Methodology, Formal analysis, Writing – original draft. Tom F. Turner: Conceptualization, Supervision, Writing – review & editing. Melanie E. Moses: Conceptualization, Writing – review & editing. Alex J. Webster: Conceptualization, Methodology, Formal analysis, Writing – review & editing.

## Declaration of competing interests

The authors declare that they have no known competing financial interests or personal relationships that could have influenced the work reported in this paper.

## Data availability

All data are publicly available as cited in the methods.

## Code availability

The computer code associated with this project is available at the following repository: https://github.com/elizagilbert/RiverDryingDrivers

## Acknowledgements

We would like to thank Trevor Birt and Dr. Grace Haggerty of the New Mexico Interstate Stream Commission for proposing use of the Rio Grande River Eyes Monitoring Program data to better understand factors contributing to flow intermittency. We would also like to thank Chad McKenna of GeoSystems Analysis Inc. who curates the River Eyes Monitoring Program data and provided helpful details on the history of its acquisition.

## Declaration of generative AI and AI-assisted technologies in the writing process

During the preparation of this work the author(s) used ChatGPT in order to clarify sentences or paragraphs and as a way to break through writers block. After using this tool/service, the author(s) reviewed and edited the content as needed and take(s) full responsibility for the content of the publication.

## Supplemental Information

**Table S1.** Estimated temperature and precipitation parameters (mean and 95% confidence interval) for two-, three-, and ten-subreach models predicting flow intermittency. The 95% confidence interval are shown in parenthesis.

| Model | Temperature | Precipitation |
| --- | --- | --- |
| Two subreaches | 0.034 (0.026, 0.041) | -0.009 (-0.015, -0.003) |
| Three subreaches | 0.024 (0.018, 0.029) | -0.008 (-0.011, -0.005) |
| Ten subreaches | 0.029 (0.028, 0.030) | -0.011 (-0.011, -0.011) |

**Fig. S1.**
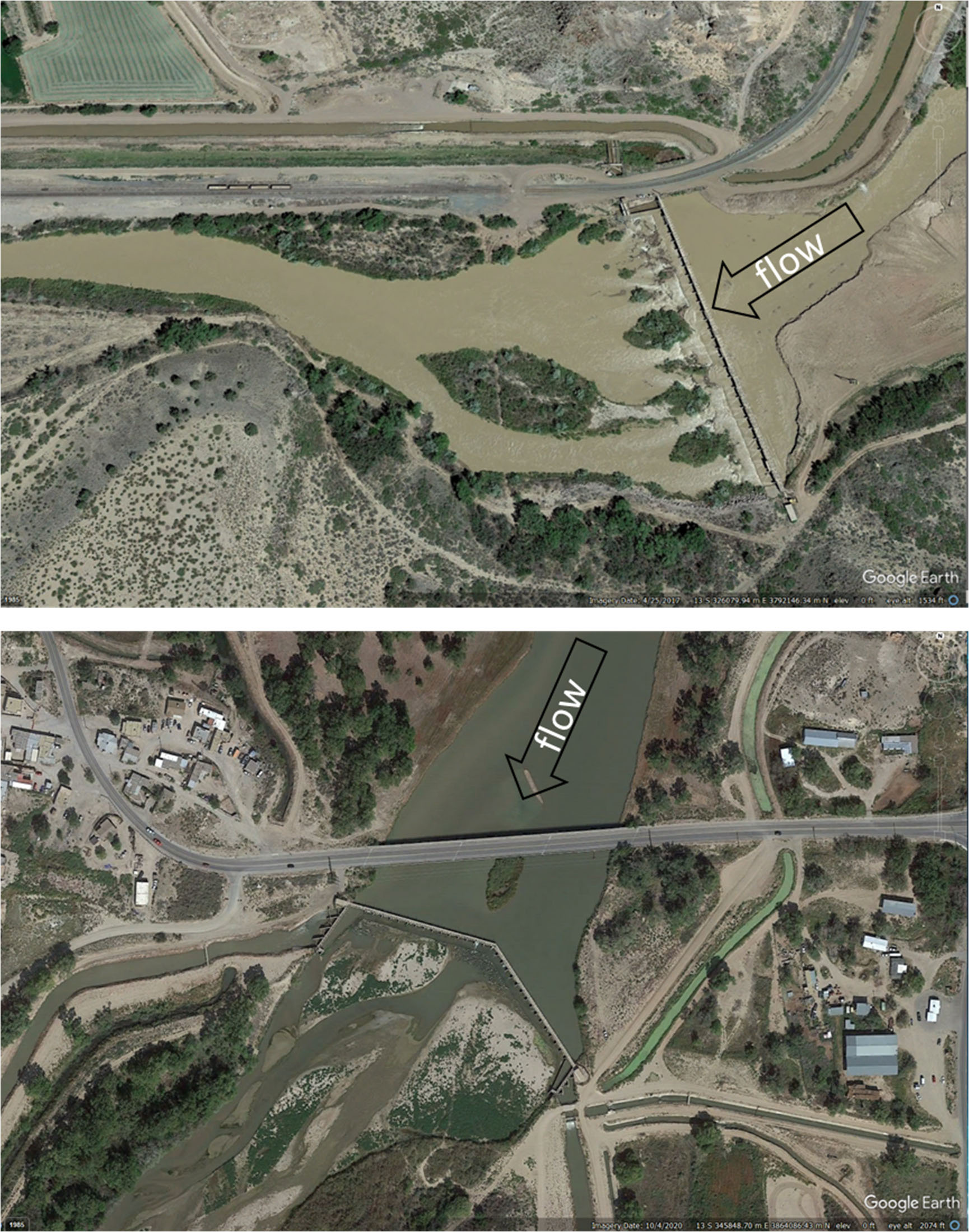
The two cross-river irrigation diversion dams present in our study area. The top picture is the most upstream diversion dam (UTM: 13 S 345322 m E 3864129) and the bottom is the diversion dam present mid-way through the study area (UTM: 13 S 326079 m E 3792146).

**Fig. S2.**
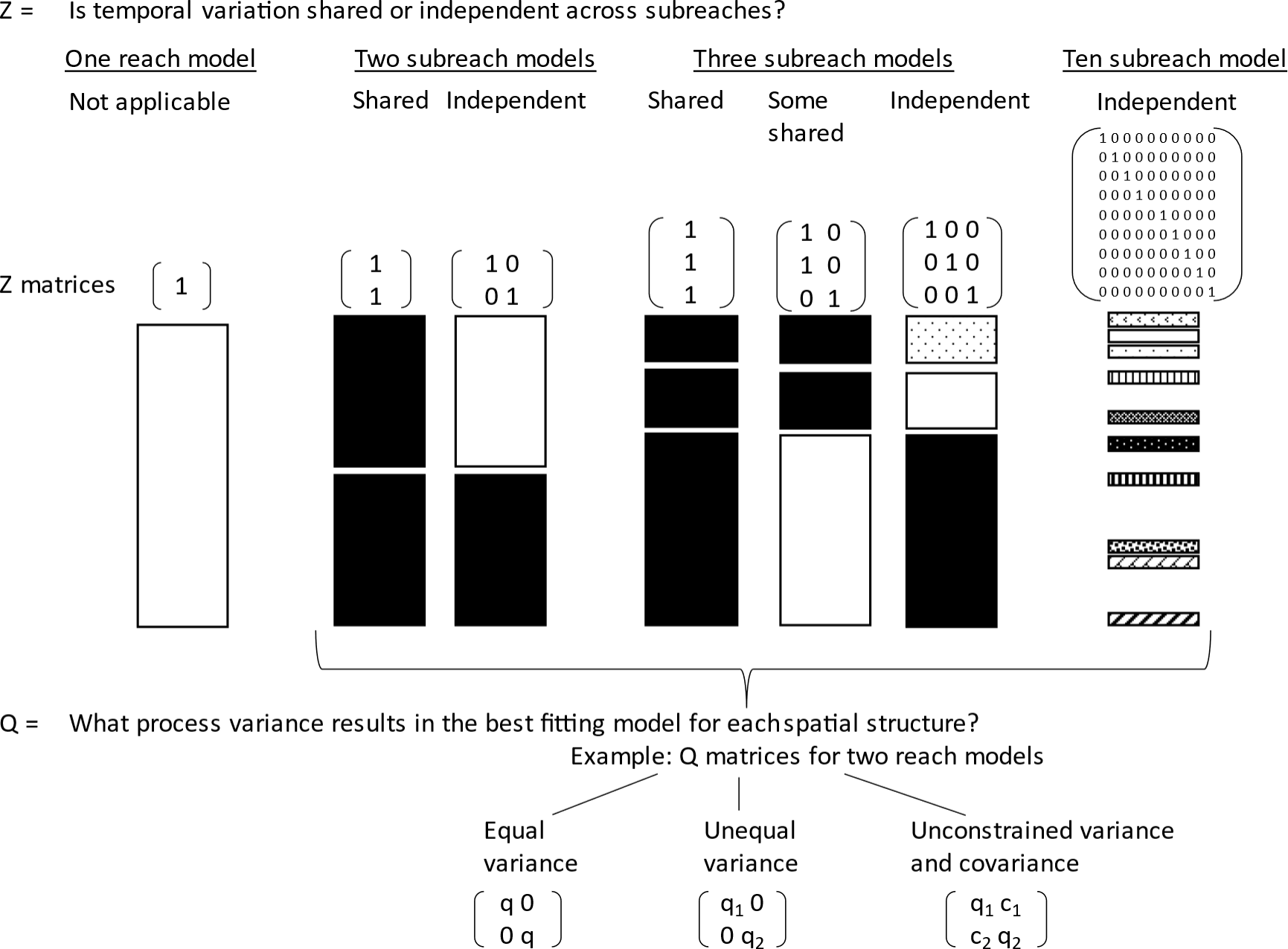
Conceptual diagram of Z and Q multivariate autoregressive state-space matrices used to analyze daily flow intermittency spatial structuring. Rectangles with the same color indicate process states whose temporal variation is shared across subreaches. In contrast, rectangles of different colors indicate process states whose temporal variation is modeled as independent.

## Notes

### Competing Interest Statement

The authors have declared no competing interest.

